# Flagellin is essential for initial attachment to mucosal surfaces *by Clostridioides difficile*

**DOI:** 10.1101/2023.05.19.541533

**Authors:** Ben Sidner, Armando Lerma, Baishakhi Biswas, Leslie A. Ronish, Hugh McCullough, Jennifer M. Auchtung, Kurt H. Piepenbrink

## Abstract

Mucins are glycoproteins which can be found in host cell membranes and as a gelatinous surface formed from secreted mucins. Mucosal surfaces in mammals form a barrier to invasive microbes, particularly bacteria, but are a point of attachment for others. *Clostridioides difficile* is anaerobic bacterium which colonizes the mammalian GI tract and is a common cause of acute GI inflammation leading to a variety of negative outcomes. Although *C. difficile* toxicity stems from secreted toxins, colonization is a prerequisite for *C. difficile* disease. While *C. difficile* is known to associate with the mucus layer and underlying epithelium, the mechanisms underlying these interactions that facilitate colonization are less well-understood. To understand the molecular mechanisms by which *C. difficile* interacts with mucins, we used *ex vivo* mucosal surfaces to test the ability of *C. difficile* to bind to mucins from different mammalian tissues. We found significant differences in *C. difficile* adhesion based upon the source of mucins, with highest levels of binding observed to mucins purified from the human colonic adenocarcinoma line LS174T and lowest levels of binding to porcine gastric mucin. We also observed that defects in adhesion by mutants deficient in flagella, but not type IV pili. These results imply that interactions between host mucins and *C. difficile* flagella facilitate the initial host attachment of *C. difficile* to host cells and secreted mucus.

## Introduction

*Clostridioides difficile* (formerly *Clostridium difficile* ^*1*^) is an opportunistic, spore-forming pathogen capable of causing gastrointestinal illness. Disease severity ranges from mild, self-limiting diarrhea to life-threatening pseudomembranous colitis or toxic megacolon. Recent emergence of hypervirulent clinical isolates with increased antibiotic resistance and rates of disease recurrence has led the United States Centers for Disease Control and Prevention (CDC) to classify *C. difficile* under the ‘urgent’ public health threat level ^*2*^. Even so, rates of community transmission continue to rise ^*3*^.

As an obligate anaerobe, vegetative *C. difficile* cells are unable to survive the aerobic environment outside the host ^*4*^. Fecal-oral transmission of dormant spores between hosts is therefore a key aspect of *C. difficile* pathogenesis. Following ingestion by the host, *C. difficile* spores transit the gastrointestinal tract (GIT), experiencing no apparent loss in viability after exposure to digestive enzymes and low pH ^*5*^. Once spores germinate, subsequent colonization of the host digestive tract is dependent upon multiple factors, including the composition and functionality of the host gut microbiome; depletion of the gut microbiota through the use of antimicrobials is one of the primary risk factors for *C. difficile* disease ^*6*^.

Competition with the gut microbiome for dietary and host-derived carbohydrates in the colon is one mechanism thought to provide protection against CDI. It was recently shown that *C. difficile* is able to cross-feed and catabolize mucin-derived glycans following enzymatic liberation by other gut bacteria ^*7*^. By controlling access to host-derived nutrients, the abundance of primary and secondary mucin degrading species may modulate *C. difficile*’s ability to colonize and proliferate. Indirectly, microbes can also influence host immunity, production of antimicrobial peptides, and mucus production, which collectively provide a barrier to colonization by *C. difficile* ^*8*^.

It has been widely demonstrated that adherence to host tissues is a prerequisite for colonization and expression of virulence factors for many enteric pathogens. For example, adherence to the host mucosa is considered to be essential for infection and disease caused by *Helicobacter pylori* ^*9*^ and *Campylobacter jejuni* ^*10*^. An early study investigating colonization of hamsters by *C. difficile* found that a virulent, toxigenic strain had greater adherence to the gut mucosa than both a less virulent toxigenic strain and a non-toxigenic strain ^*11*^. These findings provided preliminary evidence that adherence might determine virulence, but a lack of strain diversity limited formation of such broad conclusions. Later work by another group compared the adherence of 12 different isolates of *C. difficile* in a murine model of infection, finding that toxigenic strains had greater *in vivo* mucosal adherence than non-toxigenic strains ^*12*^.

In addition to receptors on the surface of epithelial cells, the luminally exposed protective outer mucus layer and extracellular matrix (ECM) offer potential receptor binding sites for intestinal microbes. Recognizing that mucosal adherence may be a determinant for virulence, Karjalainen and colleagues were interested in identifying a possible bacterial adhesin which could mediate attachment ^*13*^. A surface protein found to be upregulated following heat-shock and hypothesized to mediate adherence was recombinantly expressed and purified. The 27-kDa hypothetical adhesin adhered to Caco-2 and Vero cells, but this interaction was diminished by co-incubation with axenic murine mucus or N-acetylgalactosamine, a mucus glycan. This was one of the earliest examples of indirect evidence suggesting that *C. difficile* possesses surface proteins which bind mucus or its glycan components. Despite the significance of these findings, the relevance of possible *C. difficile* adherence to mucus during CDI is largely unknown as many investigations have focused on adherence to epithelial cells. Studies from the last decade have renewed interest in microbe-mucus interactions during CDI.

The mucus layer has unique physical and chemical properties that cause it to compress rapidly under desiccating *ex vivo* conditions such as exposure of excised tissue sections to the open air. Additionally, fixation of tissue sections by conventional cross-linking agents causes complete mucus layer compression, limiting our understanding of its architecture and microbial composition ^*14*^. However, tissue fixation with Carnoy’s fixative or paraformaldehyde has allowed for better preservation of mucus structure ^*15, 16*^ and microscopic observation of bacterial species localized to this region of the mucosa. *C. difficile* has also been found to associate with the loose outer mucus layer of the cecum and colon in a murine model of infection ^*17*^. In contrast to work from other groups, Semenyuk and colleagues found no evidence of colonization or attachment to epithelial cells, suggesting preferential colonization of the murine mucus layer by *C. difficile*.

Although ethical conflicts prohibit researchers from directly evaluating *C. difficile*’s possible association with mucus in the human gut, recent studies have provided compelling evidence that microbe-mucus interactions, including attachment, are relevant to human CDI. For example, CDI patients were reported to have decreased MUC2 mucin expression relative to healthy subjects ^*18*^. MUC2 is the most abundant mucin glycoprotein in the human colon ^*14*^ and contains many receptors important for colonization by mucus-associated bacterial species. Altered mucin expression in CDI patients could therefore plausibly impact availability of binding sites for *C. difficile*, greatly influencing virulence and disease severity.

In the same study by Engevik and colleagues, *C. difficile* alone was capable of decreasing MUC2 expression in an infection model using human intestinal organoids ^*18*^. Additionally, using mucus extracted from CDI and healthy patient stools, *C. difficile* was found to preferentially adhere to CDI patient mucus ^*18*^. A later study also reported preferential *in vitro* adherence to mucus from mucus-secreting human intestinal epithetical cell lines ^*7*^. Observed differences in mucin gene expression and relative *C. difficile* adhesion between healthy and infected populations suggest that further investigation of microbe-mucus interactions may elucidate novel targets for therapeutic development in the treatment of CDI.

Here, we probed the ability of specific mutations to *C. difficile* surface structures to impact adhesion to *ex-vivo* mucosal surfaces derived from both human and porcine sources to help elucidate the contribution of *C. difficile* interactions with host mucins to colonization. We observed that *C. difficile* bound to purified mucins *ex vivo*, that significant differences in adhesion were observed between different sources of mucin, and that flagella was important for mediating initial mucus attachment.

## Materials and Methods

### Bacterial strains and growth conditions

*C. difficile* strains 630, R20291, R20291 *pilA*, R20291 *fliC*, R20291 *pilA* p84151-*pilA* and R20291 *fliC* p84151-*fliC* were a generous gift from Dr. Glen Armstrong’s laboratory at the University of Calgary ^*19, 20*^. Based on whole-genome sequencing (Microbial Genome Sequencing Center, LLC), this *C. difficile* R20291 strain is identical to CRG3661 a strain obtained from Novartis as described by Monteford *et al*.. ^*21*^. CD2015 and CD1015 are previously characterized ribotype 027 (CD2015) and ribotype 078 (CD1015) isolates ^*22, 23*^. VPI-10463 is a previously described toxigenic isolate ^*24*^. *C. difficile*, regardless of strain, was grown in an anaerobic chamber (Coy Laboratory Products) with 5% CO- /5%H_2_/90%N_2_ atmosphere in supplemented Brain Heart Infusion medium (BHIS) using standard methods ^*25*^.

### Partial purification of commercial porcine stomach mucin

Commercial porcine stomach mucin (Sigma #M1778) was partially purified prior to use ^*26, 27*^. Crude mucin was suspended at approximately 20 mg/mL in 0.1M NaCl. The pH was adjusted to 7.0 and the solution was stirred overnight at 4°C. The following day, insoluble residues were pelleted by centrifugation (10,000 x g for 10 min.). Mucin contained in the supernatant was precipitated in ice cold 60% ethanol (v/v), then recovered by centrifugation (10,000 x g for 10 min.). Dissolving and precipitation steps were repeated twice more. The final pellet was resuspended in 0.1 M NaCl pH 7.0 and stored at -80°C. Where appropriate, O-linked glycans were removed by reductive beta-elimination ^*28*^; specifically, partially purified mucins were incubated in 0.5 M NaBH_4_ 50 mM KOH at 50°C overnight. Following glycan removal, mucin proteins were precipitated in 60% (v/v) ice cold ethanol, recovered by centrifugation (10,000 x g for 10 min.), and used to prepare mucin-coated coverslips as described above.

### Extraction and purification of porcine colonic mucin

Porcine colonic mucin was purified from the digestive tracts of adult pigs obtained from the University of Nebraska-Lincoln Department of Animal Science. Colonic mucosal tissue samples were prepared by gentle scraping of the mucosa with glass coverslips. Samples were stored at -80°C until further processing.

Extraction and purification of mucins was performed similar to previously described methods ^*29, 30*^. Briefly, mucosal tissue samples were suspended in extraction buffer (6 M guanidium chloride, 5 mM EDTA, 0.01 M NaH_2_PO_4_, 100 mM PMSF, pH 6.5) and homogenized, then stirred overnight at 4°C. The insoluble fraction containing mucins was removed by centrifugation using a JA-20 rotor (18,000 rpm, 10°C, 30 min.). The resulting pellet was resuspended in extraction buffer and the process was repeated 3 times, limiting stir durations to 2-3 hours for remaining extractions.

Extracted insoluble mucins were solubilized in reduction buffer (6 M guanidium chloride, 0.1 M Tris, 5 mM EDTA, 25 mM dithiothreitol, pH 8.0) by stirring at 37°C for approximately 5 hours. Alkylation was performed by addition of 62.5 mM iodoacetamide (final concentration) and the solution was stirred overnight at room temperature in the dark. The next day, contaminant proteins were removed by centrifugation (10,000 rpm, 4°C, 30 min.) and the resulting supernatant containing soluble mucins was lyophilized.

### Expression, extraction and purification of mucins from human epithelial cell culture lines

LS174T (ATCC CL-188) and HT29-MTX-E12 (Sigma #12040401) were grown in Dulbecco’s modified Eagle’s medium (DMEM) with 2.5 mM L-glutamine (Glutamax) and 10% fetal bovine serum (FBS) using a semi-wet interface culture method with mechanical stimulation as previously described (insert reference 29). Following passage, 10 µM DAPT (STEMCELL Technologies) was added to cells cultured in 150mm tissue culture flasks containing 30 mL DMEM + 10% FBS + Glutamax and incubated for 3-4 days in a 37°C incubator with 5% CO_2_ atmosphere until confluent to induce differentiation of goblet cells. After confluence was reached, media was replaced with 15 mL fresh DMEM and incubated with gentle rocking in a 37°C incubator with 5% CO_2_ atmosphere for three additional days. Supernatants containing mucins were collected and incubated with a mixture of protease inhibitors (1 mM PMSF, 2 mM iodacetamide) in 0.05% sodium azide and 10 mM EDTA at 4°C for 1 hour with shaking. Mucins were precipitated through the addition of 200 proof ice cold ethanol to 60% final concentration, followed by centrifugation at 15,000 X g for 30 minutes at 4°C. Pellets were washed with 66% ethanol, resuspended in filtered, deionized water containing Roche proteinase complete inhibitor, and stored at -80°C until lyophilized. Lyophized mucins were weighed to determine yield and stored at -80°C until use.

### Preparation of artificial mucosal surface

Mucins purified as described above were covalently immobilized on glass coverslips as previously described ^*31, 32*^. Briefly, coverslips were incubated in 1 M HCl overnight at 60°C, washed in sterile deionized water, then autoclaved. Once dry, coverslips were transferred to a 4% solution of (3-aminopropyl) triethoxysilane (Sigma #440140) in acetone for one hour, washed with acetone (Fisher), then heat treated at 110°C for one hour. Terminal aldehydes were formed on the surface by incubating the coverslips in 2.5% aqueous glutaraldehyde (v/v) (Fisher) for one hour, followed by rinsing with water. Finally, coverslips were incubated in 1 mg/mL mucin in phosphate buffered saline (50mM phosphate 0.15M NaCl, 10mM EDTA, pH 7.2) overnight at 4°C. Coverslips were air-dried and UV-treated for sterility, then stored at -20°C. Successful deposition of mucosal hydrogels was verified using Atomic Force Microscopy (AFM); coverslips were imaged using a Digital Instruments EnviroScope Atomic Force Microscope (ESCOPE) with a NanoScope V control station.

### *Fluorescent labeling of* C. difficile

For microscopy experiments, *C. difficile* cells were fluorescently labeled with CFDA-SE (5(6)-carboxyfluorescein diacetate succinimidyl ester; STEMCELL Technologies) ^*31*^. Briefly, mid-exponential phase cultures were washed three times with anaerobic PBS by gentle centrifugation (4,500 x g for 6 min.). After washing, cells were incubated in anaerobic PBS containing 10 mM CFDA-SE at 37°C for one hour. Following incubation, excess dye was removed by three consecutive washing and centrifugation steps as previously performed. Suspensions of labeled cells were added to sterile 6-well cell culture plates containing mucin-coated coverslips or clean glass controls and incubated at 37°C for one hour. Following incubation, unadhered bacteria were removed by washing 3 times with 1x PBS. Imaging of adhered bacteria was performed on an EVOS m5000 20x green-fluorescence objective and through Confocal Laser Scanning Microscopy (CLSM) using a Nikon A1 Confocal System with an upright Nikon Ni-E fluorescent scope and Nikon NIS-Elements using the 488 laser line with a PlanApo 60XA/1.2 WI 0.15-.18 WD water immersion lens. Adhered bacteria were quantified using CellProfiler (The Broad Institute) ^*33, 34*^.

### Mucosal adhesion assay

Prepared mucin-coated coverslips were used for evaluating the ability of *C. difficile* to interact with an artificial mucosal surface *in vitro*. Individual colonies were used to inoculate BHIS broth with 0.1% taurocholate (BHIS(TA)), and bacteria were grown to late log phase (4-6 hours) at 37°C. Cells were gently centrifuged (4,500 x g for 8 minutes), then resuspended in phosphate-buffered saline (PBS; 137 mM NaCl, 2.7 mM KCl, 10 mM Na_2_HPO_4_, 1.8 mM KH_2_PO_4_, pH 7.4) at OD600 =2.0. Suspensions were added to sterile 6-well cell culture plates containing mucin-coated coverslips or clean glass controls and incubated at 37°C for one or 24 hours. Following incubation, unadhered bacteria were removed by washing 3 times with 1x PBS. Adhered bacteria were then removed by incubation with 0.25% trypsin-EDTA solution (Gibco) at 37°C for 8-10 minutes. Trypsin was neutralized by addition of 2 volumes BHIS(TA), then recovered bacteria were plated on BHIS(TA) agar plates and enumerated after 24-48 hours of incubation at 37°C.

### Mucin binding assay in 96-well plates

In preliminary studies, 1 mg/mL partially purified porcine gastric mucin (PGM) was incubated overnight at 4°C in a sterile high-binding 96-well plate (Corning Costar REF# 3361). Wells containing 1 mg/mL bovine serum albumin (BSA) or buffer alone were added as controls for non-specific binding. The next day, wells were washed with buffer and an adherence assay involving incubation of bacteria in treated wells, washing to remove unadhered bacteria, and enumeration of adhered bacteria was performed (similar to the steps described for the mucosal adhesion assay). The relative adherence of *C. difficile* R20291 to wells treated with BSA or buffer alone was greater than to wells treated with PGM (Supplemental Figure 1).

### Swimming motility assay

To assess swimming motility, *C. difficile* strains R20291, 630 VPI 10463, CD2015 and CD1015 were cultured overnight in BHIS(TA) broth. Overnight broth cultures were used to inoculate 0.5x BHIS with 0.3% agar plates and measurements were recorded every 24 hours for 72 hours ^*35*^.

### Statistical analyses

All statistical analyses were performed using GraphPad Prism version 9.3.1 for macOS (GraphPad Software, San Diego, California USA). All significance differences were determined at p-value<0.05. Relevant statistical tests used for comparisons are discussed in figure captions.

## Results

### C. difficile *adheres to* ex-vivo *hydrogels*

Building upon previous studies measuring *ex-vivo* mucosal adhesion by microorganisms (refs), we created mucosal hydrogels from mucus samples secreted in cell-culture and from animal sources, purchased commercially or harvested directly. Figure 1 shows the process for creation of these ex-vivo hydrogels (panel A) as well as validation of mucosal deposition by atomic force microscopy (panel B). Specific adhesion was observed in panel C for *C. difficile* labeled with CFDA-SE to coverslips coated with porcine gastric mucin (PGM), with similar results obtained from confocal laser scanning microscopy (CLSM) and enumeration of adhered bacteria through colony-formation.

**Figure 1:**
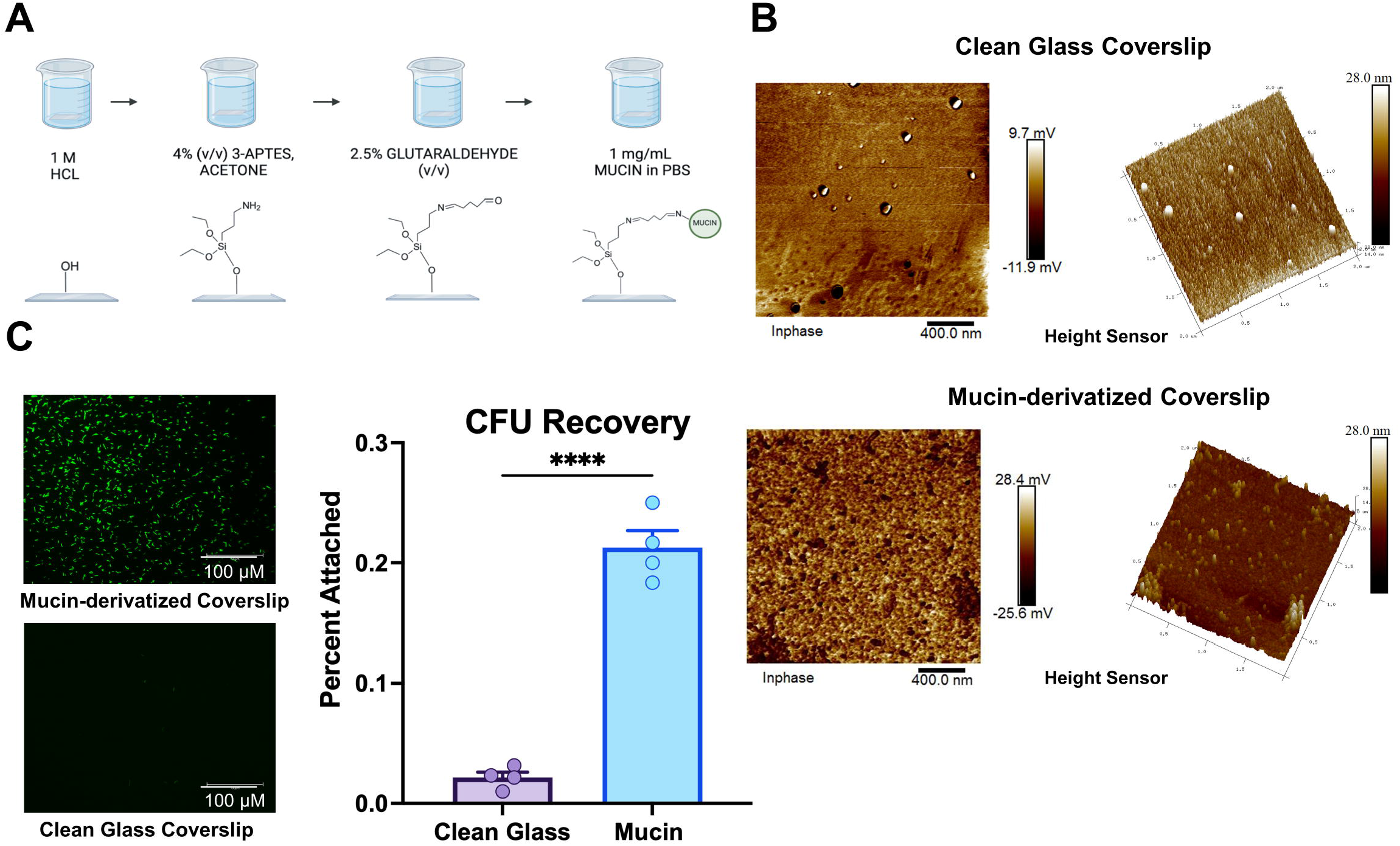
Preparation of *ex-vivo* mucosal hydrogels. A) Schematic of preparing glass coverslips with mucosal hydrogels. B) Atomic Force Microscopy (AFM) images of glass coverslips before (above) and after (below) derivatization with mucins. C) representative images from Confocal Laser Scanning Microscopy (CLSM) of *C. difficile* bacteria, derivatized with CFDA-SE, adhered to coverslips as well as quantification by CellProfiler. Significant differences were calculated using Student’s one-tailed T-test, **** p > 0.0001

### *Defects in adhesion are observed for an R20291* fliC *mutant, but not* pilA *mutant*

To probe the molecular basis for *C. difficile*-mucin interactions, we tested the binding of gene-interruption mutants of the major subunits of flagella (*fliC*) and the type IV pilus (*pilA1*) (see Figure 2, panel D). Both flagella and type IV pili have been implicated as contributing to host cell adhesion by *C. difficile* ^*36, 37*^ and other bacterial species ^*38-43*^. In this model of mucin-adhesion, we found a pronounced defect after 1 hour for the *fliC* mutant, but no defect for the *pilA* mutant, which showed an increase in binding similar to that observed by McKee ^*37*^. Unlike McKee’s cell-culture binding experiments, we saw no reversal of the *pilA1* defect after 24 hours (panel B) which may be attributable to the binding medium (PBS) which does not allow for biofilm formation.

**Figure 2:**
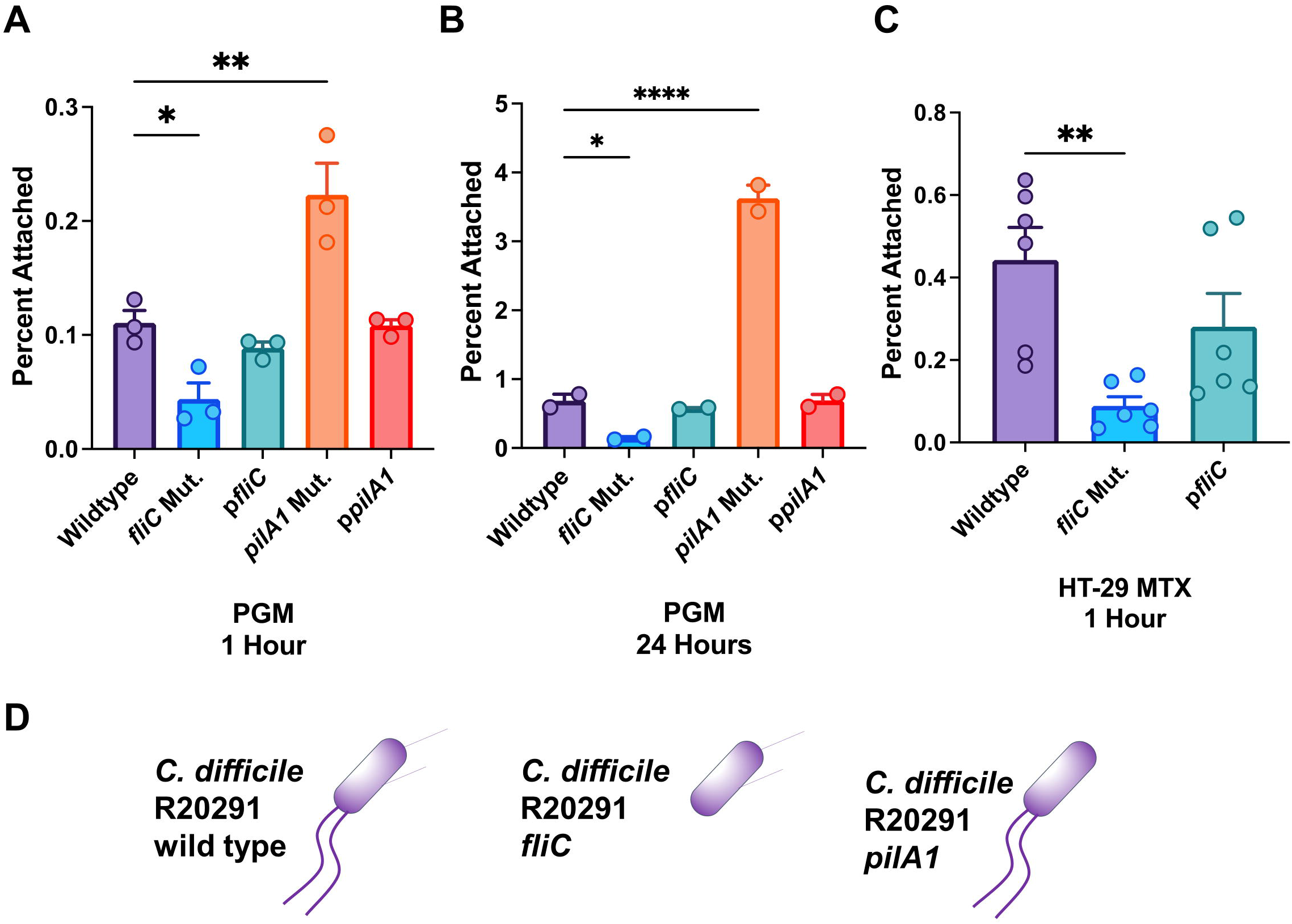
Adhesion by pilus and flagellar mutants. A-C) Adherence of *C. difficile* R20291 and gene-interruption mutants to *ex-vivo* mucosal hydrogels of A) Porcine Gastric Mucin (PGM) (1 hour), PGM (24 hours) and C) mucus secreted by HT-29 MTX cells (1 hour). D) Schematic showing surface appendages of *C. difficile* R20291 and mutants. Significant differences are measured from the wild type strain using Student’s one-tailed T-test, * p > 0.05, ** p > 0.01, **** p > 0.0001

To verify that this defect was not specific to porcine gastric mucins, we also measured adhesion to hydrogels derived from mucins secreted by HT29-MTX-E12 cells. We found that these surfaces showed an identical defect in binding for the *fliC* mutant. This defect is consistent with the binding defect observed for an R20291 *fliC* mutant by Baban *et al*. ^*36*^.

### *Flagellar motility is not a prerequisite for* C. difficile *mucin adhesion*

Because flagella could contribute to attachment to surfaces both through swimming motility and direct adhesion, we used a panel of *C. difficile* strains, both motile and non-motile as measured by swimming motility, for their adhesion to *ex-vivo* mucosal hydrogels. Figure 3 shows the results of mucin adhesion (panel A) and swimming motility (panel B) assays for a panel of five *C. difficile* strains, CD1015, VPI 10463, CD2015, R20291 and CD630. The two 027 ribotype strains (CD2015 and R20291) show the highest swimming motility, but only moderate adhesion to a PGM surface. VPI 10463 and CD630 both show weak motility and adhesion, while CD1015 (078 ribotype) shows the strongest adhesion but little or no swimming motility. Based on these results, we conclude that robust attachment to mucus by *C. difficile* in this assay is possible even in the complete absence of flagellar motility.

**Figure 3:**
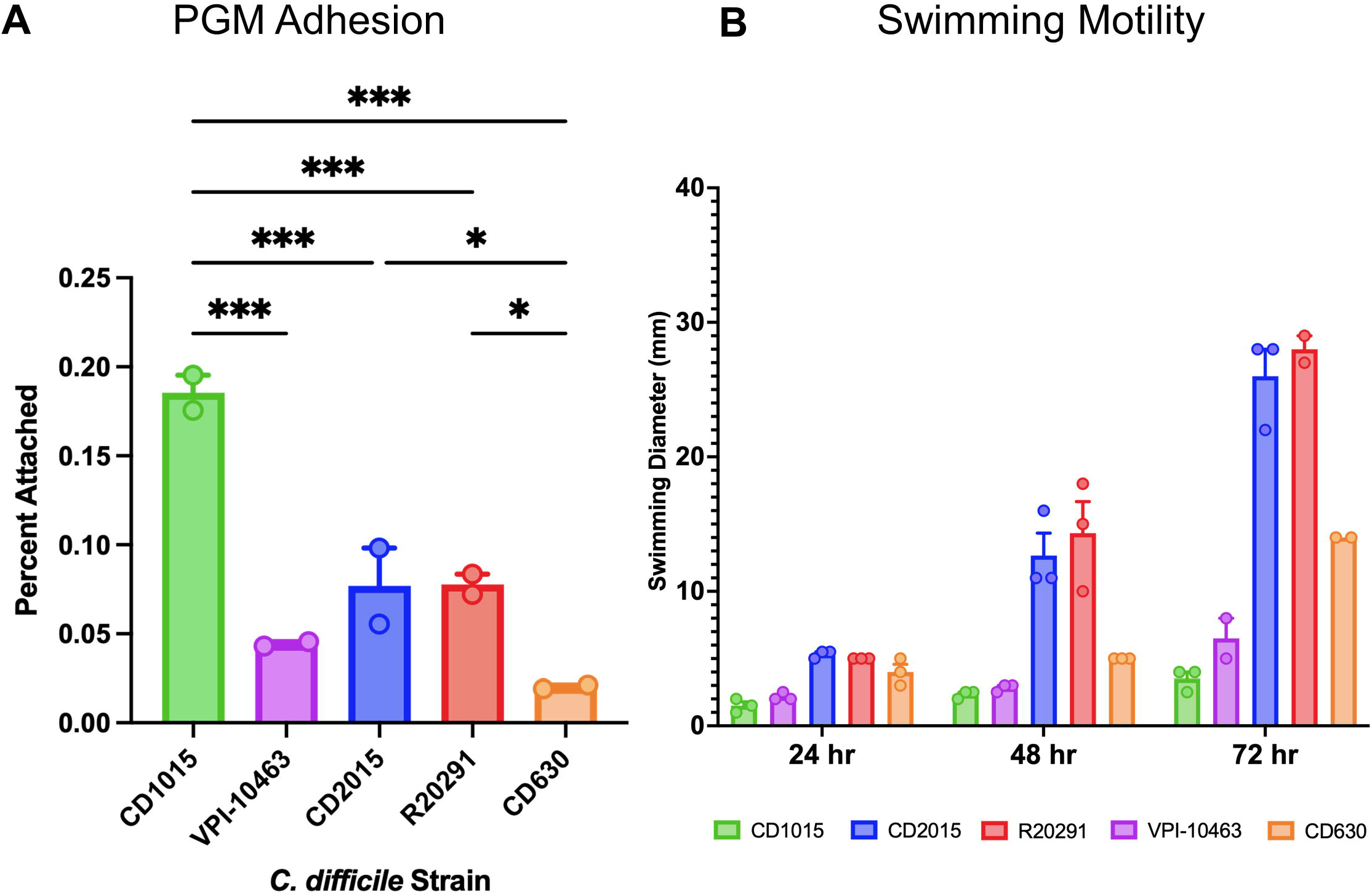
Motility and Adhesion of *C. difficile* strains. A) Adherence of *C. difficile* strains to PGM. B) Swimming motility in 0.2% agar, measured as the diameter of the bacterial expansion after 24, 48 and 72 hours. Using Student’s one-tailed T-test, all differences between groups were calculated with only significant differences shown for panel A, * p > 0.05, ** p > 0.01, *** p > 0.001. In panel B, at 72 hours, all differences are significant except the pairs R20291/CD2015 and CD1015/VPI-10463.

### *Mucin composition and glycosylation impact* C. difficile *adherence*

Mucus is a hydrogel composed of various mucin glycoproteins forming a heterogenous gelatinous matrix. The composition varies depending on both the identity of the mucin polypeptides (for example, MUC1, MUC2, or MUC5AC) and the pattern of glycosylation, which is likely to be heterogenous even for the individual mucin proteins. The degree of glycosylation of naturally-occurring mucins is such that all interactions with mucins are typically presumed to occur through the glycans, rather than the polypeptide itself ^*44*^ and both the protein composition and glycosylation pattern vary based on the source of the mucus in a species and site-specific manner ^*45*^. To probe the specificity of *C. difficile* adhesion for distinct, naturally-occurring mucosal compositions, we tested the adhesion of *C. difficile* R20291 to mucosal hydrogels from porcine gastric and colonic mucins as well as two human colonic cell culture lines (HT29-MTX-E12 and LS174T). Figure 4 panel A shows the results of mucosal adhesion assays performed with and without a reduction in O-glycosylation through β-elimination. As expected, the removal of O-glycans significantly reduced adhesion by *C. difficile*. The residual binding observed could be attributed to direct interactions between *C. difficile* and the mucin polypeptides, binding to N-linked glycans. We hypothesize instead that the β-elimination was incomplete, with the remaining O-linked glycans being capable of binding.

**Figure 4:**
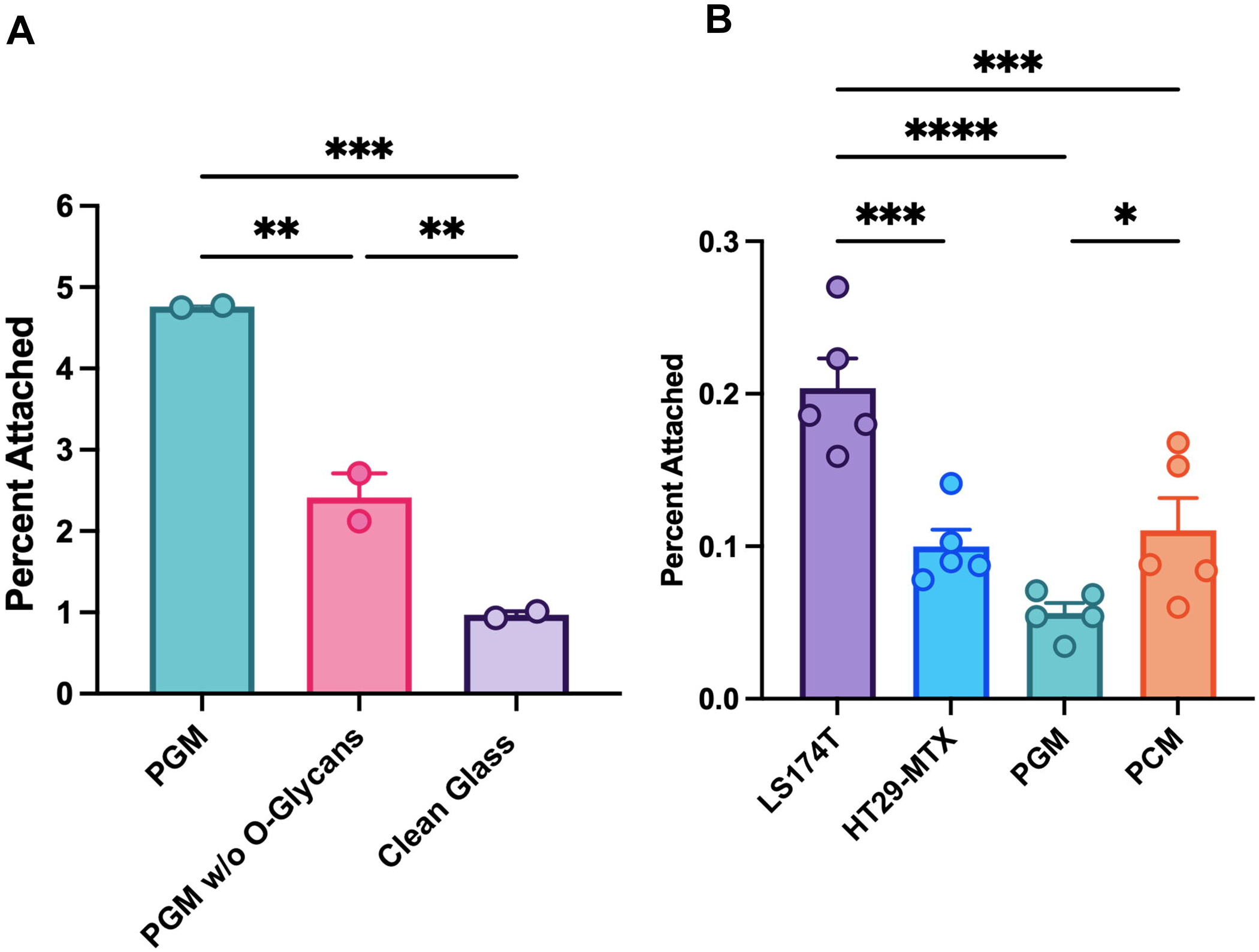
Effects of mucosal composition on *C. difficile* adhesion. A) Adherent bacteria recovered from PGM, with and without β-elimination to remove O-glycans and to non-derivatized glass coverslips. B) Adherence by *C. difficile* R20291 to mucosal hydrogels from various sources. Significant differences between all groups were calculated using Student’s one-tailed T-test * p > 0.05, ** p > 0.01, *** p > 0.001, **** p > 0.0001

Comparing adhesion to mucosal hydrogels from different mucus sources (Figure 4, panel B), we observed that *C. difficile* R20291 adhered to porcine colonic mucins at a significantly higher level than porcine gastric mucins, consistent with the hypothesis that *C. difficile* is adapted to colonization of the GI tract and does maintain the capacity bind the sulfated glycans of gastric mucins ^*46*^. We also found significantly greater binding to LS174T-derived hydrogels than those derived from HT-27 MTX cell culture. This difference could be attributable to higher relative expression of MUC2 in LS174T cells ^*47*^ as MUC2 has previously been hypothesized to contain receptors for *C. difficile* adhesion ^*7*^.

## Discussion

Historically, the layers of mucus found in the lungs, GI, and nasopharyngeal tracts have been thought of as barriers to colonization by microbes ^*48*^. However, recent results from this study and others ^*7*^ show specific adhesion of a GI pathogen to reconstituted host mucosa. One question raised by these results is to what extent is the adhesion observed by *C. difficile* (and other GI bacteria) for eukaryotic cells in culture is due to adhesion to membrane bound mucins. Membrane-bound mucins are bound to the cell membrane by transmembrane (TM) α-helices, but can be released in some cases by proteolytic cleavage of the C-terminal extracellular domain ^*49-51*^.

Figure 5 shows three potential adhesive processes leading to *C. difficile* colonization; adherence to the intestinal lumen mediated by direct binding to mucins, attachment to the host epithelium, which could involve membrane-bound mucins or other surface receptors and finally, biofilm formation by *C. difficile*, which increases the number of host-adhered cells indirectly through interactions between bacterial cells. The reduction in binding for an R20291 *fliC* mutant to both Caco-2 cells (Baban et al. ^*36*^) and mucin hydrogels (this study) implies that interactions with mucins may contribute to *C. difficile* attachment to both the lumen and the host epithelium under native conditions. The relative importance of each site of attachment, and biofilm formation at either, during infection is unclear; the high turnover of intestinal mucus may make it unsuitable for long-term colonization, but it adherence to the lumen may be advantageous for an obligate anaerobe because oxygen levels are somewhat lower than the epithelial surface ^*52*^, though biofilm formation may offer some protection against ambient O_2_.

**Figure 5:**
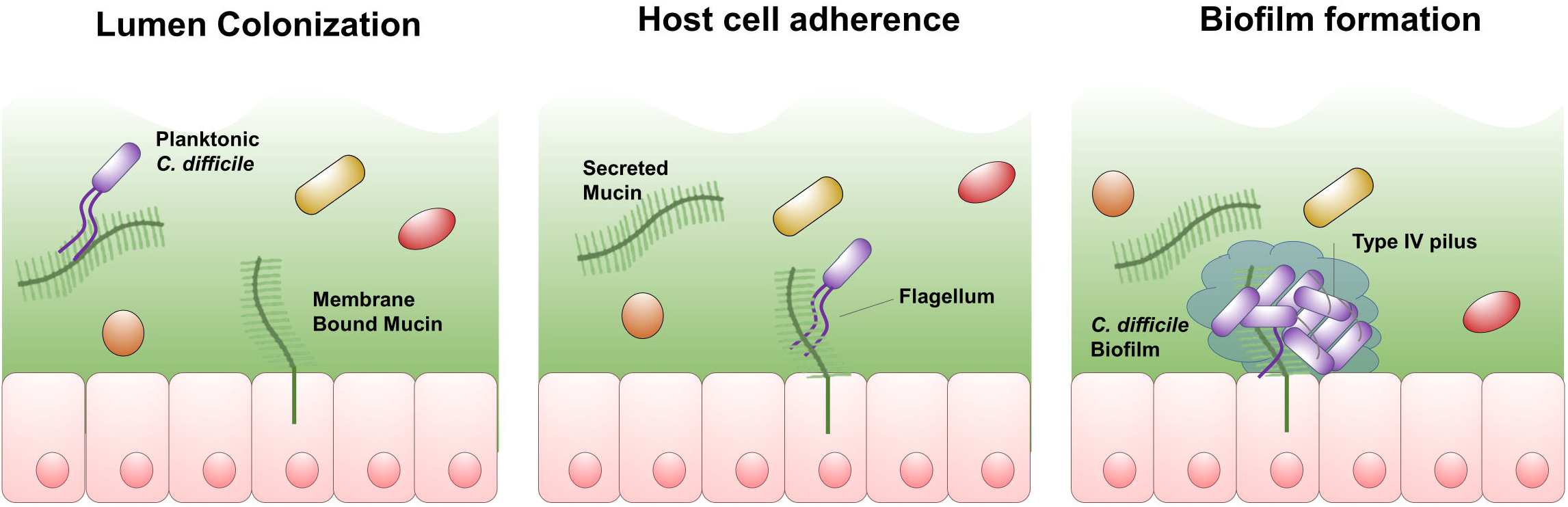
Model of *C. difficile* colonization. Schematic representation of association of *C. difficile* with the host lumen, epithelial cell layer and biofilm formation during infection.

Because type IV pili promote not only biofilm formation ^*25, 40, 53-59*^ but also host cell adhesion in a number of bacterial species ^*39, 43, 60, 61*^, the lack of an adhesion defect in the *pilA1* mutant is somewhat counter-intuitive, particularly in light of the defect seen in cell-culture and in a mouse model ^*37*^. We hypothesize that the promotion of biofilm by *C. difficile* type IV pili, as shown in Figure 5, increases association with host indirectly as has been observed previously for type IVb pilus systems such as the Bundle-Forming Pilus (BFP) of enteropathogenic *E. coli* (EPEC) ^*62, 63*^. Our group previously published the structure of PilA1 which does resemble the major subunits of type IVb pili ^*19*^.

In Figure 5 attachment to host mucins (in the lumen and on the cell surface) is depicted as occurring directly through *C. difficile* flagella, based on the results reported here, reports from other groups of flagellar adhesion by *C. difficile* ^*64, 65*^, and known interactions between FliC and host mucin glycoproteins in other infectious bacteria ^*66, 67*^. However, we did not find that the use of purified *C. difficile* flagella and flagellar proteins as competitive inhibitors resulted in significant reductions in binding using this model (Supplemental Figure 3). An alternative hypothesis is that the expression of *C. difficile* FliC is necessary for the production of downstream adhesins which adhere to host glycoproteins. Previous studies have pointed to interactions between FliC and FliW, which regulates the activity of the pleiotropic regulatory protein, CsrA; deletion of *csrA* reduces adhesion, although this could be mediated by CsrA-mediated regulation of *fliC* ^*68*^.

A related question is to what extent the mechanisms of host adhesion are conserved between *C. difficile* strains. Unlike the defect in binding observed for the R20291 *fliC* mutant for mucins here and for Caco-2 cells previously ^*36*^, Dingle *et al*. observed no defect in binding for *fliC* or *fliD* mutants of *C. difficile* 630 to Caco-2 cells ^*20*^. The robust adhesion observed for CD1015 in this study also calls into question a direct role for flagella in adhesion, as CD1015, a ribotype 078 strain, shows no swimming motility. CD1015 is missing the f3 flagellar operon and the extent to which it produces flagellar structures is unknown; bacteria which produce FliC but are missing other flagellar genes have been shown to produce truncated flagellum-like structures in other bacterial species ^*69, 70*^. Isolation of *C. difficile* surface appendages through vortexing and precipitation showed what appeared to be band for FliC in SDS-Page for CD1015 (Supplemental Figure 2B) but use of these precipitated proteins as competitive inhibitors showed no effect in our binding assays (Supplemental Figure 3B).

Based on previous studies and the results described here, we hypothesize that direct interactions between bacterial flagellar subunits and host glycoproteins contribute to colonization in a variety of bacterial species. To the extent that interactions with the host mucosa could contribute to *C. difficile* colonization, competition between microbes for host mucins as attachment sites and microbial degradation of host mucins as nutrient sources are potential factors influencing the competition between *C. difficile* and the commensal microbiome. Future studies of *C. difficile* adhesion must evaluate this hypothesis and others to provide a molecular lens on the point of attachment. Ultimately an atomistic description of the processes of attachment by this bacterial pathogen will provide opportunities for the development of specific, targeted therapeutic interventions.

## Acknowledgements

We would like thank Melinda Engevik (Medical University of South Carolina) for providing advice, protocols, and cell lines for purification of mucins from human epithelial cells, as well as Eric Martens and Sadie Gugel for providing advice and protocols for purification of mucins for porcine colonic tissue. We also extend our appreciation to Thomas Burkey, Gary Sullivan, Calvin Schrock and the staff of UNL’s Loeffel Meat Laboratory for providing access to porcine colonic tissue from deceased animals that were used to purify porcine colonic mucins. We would also like to acknowledge excellent technical support from Joe Zhou and the staff of the UNL microscopy core as well as Lanping Yue and the staff of the UNL Surface and Materials Characterization Facility.

## CRediT author statement

**Ben Sidner:** Investigation, Formal analysis, Writing - Original Draft. **Armando Lerma:** Investigation. **Baishakhi Biwas:** Investigation. **Leslie Ronish:** Validation. **Hugh McCullogh:** Methodology. **Jennifer Auchtung:** Supervision, Funding acquisition, Writing - Review & Editing. **Kurt Piepenbrink:** Supervision, Funding acquisition, Writing - Review & Editing.

