## Supporting Material for "Flagellin is essential for initial attachment to mucosal surfaces *by Clostridioides difficile*"

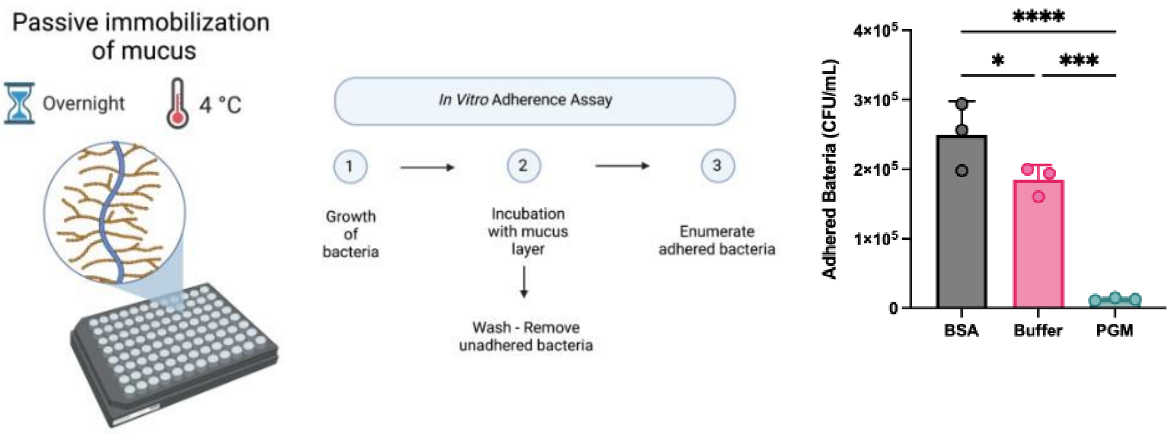


**Supplementary Figure 1**: Immobilization of mucus onto high-binding plates. Porcine Gastric Mucus (Sigma) was attached to high-binding plates overnight at 4C and blocked with bovine serum albumen (Fisher) Adherence by Cd R20291 was measured after incubation for one hour at 37C by enumeration of colony forming units after washing and removal by trypsin (See Methods). Greater adherence was measured to wells not coated with mucins (with or without blocking with BSA), suggesting that non-specific adhesion to plastic surfaces by *C. difficile* is potentially equal in strength to many biomolecular interactions. Significant differences were calculated using Student's one-tailed T-test, * $p > 0.05$, ** $p > 0.01$, *** $p > 0.001$, **** $p > 0.0001$

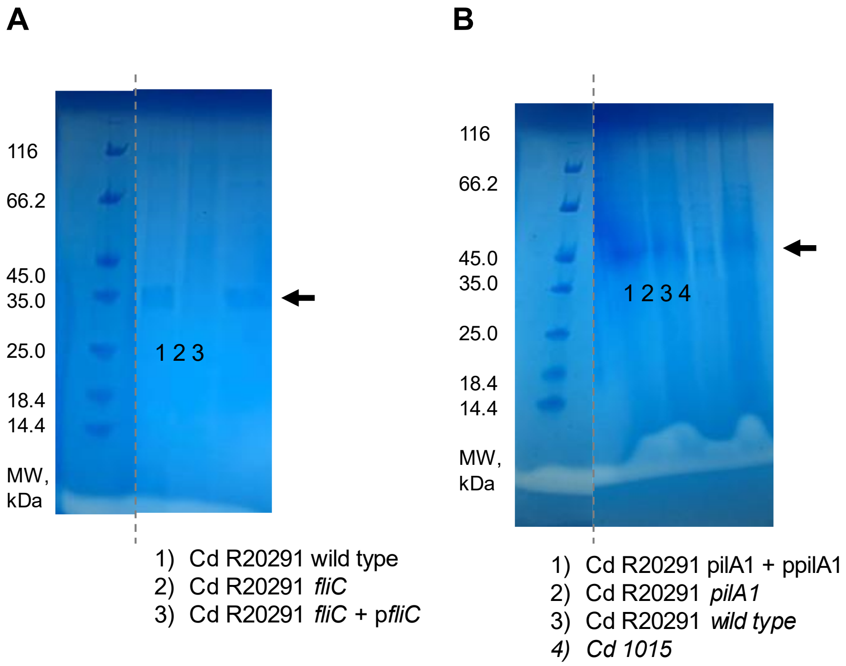


**Supplementary Figure 2**: Isolation of *C. difficile* flagella through shearing. A) After 24 hours of growth in BHIS broth, flagella were sheared from the surface of *C. difficile* R20291 and mutants through vortexing (10 seconds) alternated with incubation on ice (20 seconds) for a total of 60 seconds of vortexing in anaerobic sealed tubes. The suspension was centrifuged at 4000 x g for 10 minutes to pellet the cells. The pellets were discarded and ammonium sulfate was then added to the supernatant at 30% and they were stirred overnight at 4C. These suspensions were centrifuged at 20,000 x g for 30 minutes to pellet any precipitated proteins. Those pellets were resuspended in PBS at run on SDS page shown above. B) Identical methods were employed to recover flagella from the surface of the Cd R20291 pilA1 mutant and complement as well as Cd 1015 wild type.

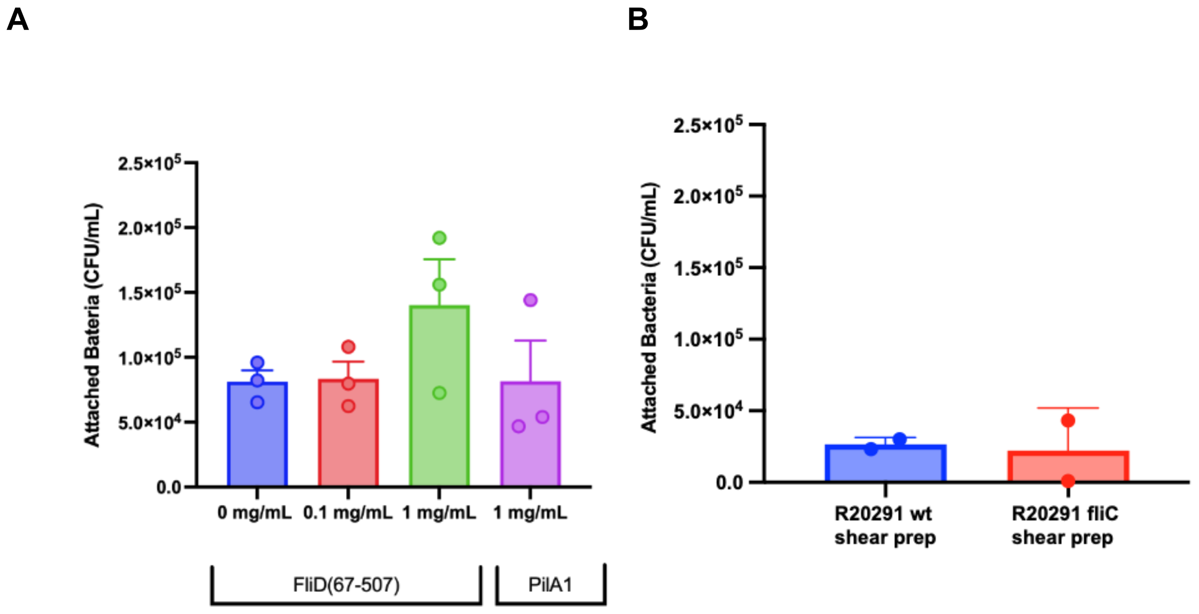


**Supplemental Figure 3**: Competitive inhibition of Cd FliD and isolated flagella in mucosal binding assays. Mucosal binding experiments with Cd R20291 binding to porcine gastric mucin were performed as previously described with the addition of either A) Cd FliD from recombinant expression in *E. coli* or B) flagella sheared from cell surface as described above at approximately 1 mg/ml. No differences between groups were significant.
